# Deforestation reduces the genetic structure of an epiphytic weed across space and time: an IBM approach

**DOI:** 10.1101/2020.10.19.345454

**Authors:** Cleber Juliano Neves Chaves, Bárbara Simões Santos Leal, Davi Rodrigo Rossatto, Uta Berger, Clarisse Palma-Silva

## Abstract

Deforestation has allowed the massive dispersal and reproduction of some plants that are commonly referred to as weeds. The rapid spread of many weeds into newly disturbed landscapes is often boosted by clonal growth and self-fertilization strategies, which conversely increases the spatial genetic structure (SGS) of populations and reduces the genetic diversity. Here, we use empirical and modeling approaches to evaluate the spatio-temporal SGS dynamics of *Tillandsia recurvata* (L.) L., a common epiphytic weed with selfing reproduction and clonal growth widespread in dry forests and anthropically deforested landscapes in North and South America. We constructed an individual-based model (IBM) and adjusted the parameters according to empirical genetic data, to simulate the spreading of *T. recurvata* over time and across random landscapes with distinct tree densities. From empirical data, we observed a strong SGS among *T. recurvata* subpopulations hosted on neighbor trees and a contemporary spread from several population sources. Our model shows that the highest SGS appear in landscapes with more than 200 trees/ha and up to the 5^th^ year of colonization of open landscapes (ca. 100 trees/ha) when SGS starts to reduce drastically. These results suggest that the deforestation commonly observed in anthropically transformed landscapes may reduce the dispersal limitation and genetic structure of *T. recurvata* subpopulations, creating suitable conditions for the rapid spread of *T. recurvata* from multiple surrounding sources. The combination of clonal growth and self-fertilization with the optimal conditions created by anthropogenic transformations may explain the spreading success of *T. recurvata* and other weeds into new landscapes. Our results indicate that the drastic reductions in tree densities induced by human-modifications in natural landscapes may lead to a partial loss of resistance for dispersal by wind and increased the conditions for *T. recurvata* to develop massive populations in anthropogenic landscapes.

## INTRODUCTION

Intensive disturbances are inherent to human history and have substantial effects on natural communities (de Wet and Harlan 1975). Ecosystems are currently subjected to unprecedented rates of human-induced changes that are fostering the sixth mass extinction (Ceballos *et al.* 2015). However, not all organisms respond the same way to human disturbances. While many species become extinct or migrate, others thrive in anthropogenic environments and can spread into natural environments (Wilcove 1985; Airoldi and Bulleri 2011). Indeed, human activities, such as those related to deforestation, have allowed the dispersal and reproduction of many species, such as those commonly referred to as weeds (Airoldi & Bulleri 2011). Weeds are opportunistic species that can evolve from introduced or native species that grow within human-transformed landscapes without being cultivated and negatively affecting both the environment and the economy (Baker 1974; de Wet and Harlan 1975; Richardson *et al.* 2008).

Between 50 and 80% of all invasive plant species can be classified as weeds, depending on current impacts and human perception (Richardson *et al.* 2008). However, native species possessing traits that pre-adapt them to establish in newly disturbed areas can also evolve into weeds (van Etten *et al.* 2017). In addition to landscape disturbances, propagule pressure, and individual density, the spreading rate of weeds is mainly driven by their high dispersal abilities and wide plasticity, which enable them to rapidly colonize large areas (Baker 1974; Booth *et al.* 2002; Kuester *et al.* 2014; van Kleunen *et al.* 2018; De Bona *et al.* 2019). In particular, clonal growth and self-fertilization (hereafter referred to as ‘selfing’) are some of the most important adaptations that allow weeds to establish new populations after long-distance dispersal events (Baker 1955; Razanajatovo *et al.* 2016). Despite their entirely different origins, both clonal and selfing abilities help increase the invasiveness of a species by mitigating the Allee effect of small populations and, therefore, enhance reproductive assurance and genetic transmission (Vallejo-Marín *et al.* 2010; Rodger *et al.* 2013; Barrett and Harder 2017). While selfing can increase species' investment in seed production, clonal growth speeds up ramet production and reduces the likelihood of genet death by sharing stochastic risk over multiple ramets (Klimeš *et al.* 1997; Barrett 2015).

Despite the positive effect of cloning and selfing strategies on weed invasiveness, the aggregation of self-fertilized offspring and clonal individuals strongly influences the spatial genetic structure (SGS) of populations, reducing local genetic diversity and promoting inbreeding depression (Heywood 1991; Barrett 2013; Barrett *et al.* 2014; Barrett 2015). In turn, clonal growth may lead to the maintenance of (once established) genetic diversity (Ellstrand and Roose 1987; Loh *et al.* 2015), while continuous selfing can result in the absence of inbreeding depression as a consequence of purging of deleterious alleles over generations (Hedrick 1994; Arunkumar *et al.* 2015; Barrett and Harder 2017). These effects lead to a strong population subdivision, creating a metapopulation dynamic with extinction in a site being balanced by recolonizations. This dynamic selects selfing and clonal genotypes, due to their higher capacities of recolonization (Pannell and Barrett 1998; Barrett 2013).

Understanding local spread dynamics of weeds is essential to elucidate how native selfing and clonal species opportunistically reach broad range distributions, despite their tendency to form highly aggregated populations with low genetic diversity. Aside from the aggregation tendency of selfing and clonal species, individuals within populations with reduced abundance and limited seed dispersal usually are highly related at the beginning of colonization (Hamrick and Trapnell 2011). With an increase in abundance after colonization, the kinship among individuals often reduces due to greater overlapping of maternal seed shadows and successive introductions of new genotypes through seed dispersal from neighbor populations (Hamrick and Trapnell 2011; Côrtes *et al.* 2013). Therefore, density-relatedness dynamics is expected to reduce the high SGS expected for populations of selfing and clonal species throughout colonization stages (Hamrick and Trapnell 2011; Trapnell *et al.* 2013; Chung *et al.* 2018).

Aside from the intrinsic effects of clonal growth and self-fertilization, the deforestation seems to be a turning point in which some natural species develop fast colonization abilities and form abundant populations into newly disturbed landscapes (Cardelús et al. 2005; Roberts et al. 2005; Quaresma et al. 2018). To shed light on this topic, here we integrate empirical and simulated genetic data to study the spatio-temporal dynamics of the local spread of *Tillandsia recurvata* (L.) L., an epiphytic weed with selfing and clonal growth strategies widespread in canopies of human-modified landscapes in the American continent (e.g. Claver et al. 1983; Bartoli et al. 1993; Flores-Palacios et al. 2015a). Specifically, we aimed (I) to investigate whether subpopulations of *T. recurvata* growing on neighbor trees of an anthropical landscape are genetically structured; (II) to examine the temporal dynamics of genetic structuring of *T*. *recurvata* subpopulations in an anthropical landscape; as well as (III) to examine whether the genetic structuring of *T. recurvata* varies according to tree densities in each landscape. Since obtaining continuous empirical genetic data over a long period can often be unfeasible (Epps and Keyghobadi 2015), modeling approaches are helpful to understand spatio-temporal effects on the genetic structure. Individual-based models (IBMs) explicitly incorporate characteristics of individuals (e.g., life stage, size, and genotype) and simulate interactions among them and with their surroundings (Grimm and Railsback 2005). Herein, we employed microsatellite markers and an individual-based model (IBM) to estimate the fine-scale SGS of a *T. recurvata* population scattered on neighboring trees and to understand the emergence of the observed empirical patterns of genetic variation over colonization time and across landscapes with distinct tree densities.

## MATERIAL AND METHODS

### Model species

*Tillandsia recurvata* is an epiphytic bromeliad distributed in the American continent. This species is known by several popular names (e.g., ball-moss, Jamaican ball moss, *musgo de bola*, *heno de bola*, *galinita*, and *cravo-do-mato*; Birge 1911), which refer to the ‘dry aspect of leaves and the intense clonal growth from leaf axils that forms a ball-like shape. Particularly found in open biomes from Argentina and Brazil to the southern United States (Smith and Downs 1977; GBIF 2017), the high abundance and dominance of *T. recurvata* in epiphytic communities constitutes a characteristic feature of landscapes and has long attracted the interest of naturalists and researchers (e.g. Birge 1911; McWilliams 1992; Flores-Palacios et al. 2014; Orozco-Ibarrola et al. 2015). Studies have described *T. recurvata* as the most xerophytic species of tropical and subtropical epiphytes (e.g., Benzing 2000; 2012), with “probably the greatest adaptability of any plant in the Western Hemisphere” (Foster 1945:14). Several ecological and reproductive features, such as small size, absorbent leaf scales, CAM photosynthesis, wind-dispersed seeds, the spontaneous self-pollination underpinning its cleistogamous flowers, and intense clonal growth have allowed *T. recurvata* to generate large populations even on isolated trees within recently disturbed landscapes (Soltis *et al.* 1987; Smith *et al.* 1989; Orozco-Ibarrola *et al.* 2015; Chilpa-Galván *et al.* 2018). Each host tree can thus act as an isolated patch of habitat scattered in a harsh matrix (Southwood and Kennedy 1983; Burns 2007), determining that the majority of dispersed seeds of *T. recurvata* fall close to the mother and establish on the same host tree (e.g., Trapnell *et al.* 2004; Victoriano-Romero et al. 2017; Torres *et al.* 2018).

The high adaptability of *T. recurvata* has fostered the species distribution in human-disturbed regions, forming large populations in urban landscapes, actively growing on fences and electric light wires (e.g., Martins 2009; Chaves et al. 2016). Such great abundance has led some authors to consider *T. recurvata* as an “epiphytic weed,” and some of them even sought to describe herbicides and other techniques to control its spreading (e.g., Claver et al. 1983; Bartoli et al. 1993). Although *T. recurvata* cannot be classified as a parasite, studies have listed deleterious effects that massive populations may have on co-occurring epiphytes and host trees, as it competes for light with new shoots, modifies barks, and affects seedling growth (e.g., Birge 1911; Flores-Palacios et al. 2014; Aguilar-Rodriguez et al. 2016). Therefore, the reproductive biology and opportunistic behavior of *T. recurvata* make this species a good model species for studying the temporal and spatial spreading dynamics of weeds.

### Empirical genetic data

The empirical study took place in an anthropically transformed landscape of ca. 0.2 ha in southwest Brazil (−21.244289° S, -48.300486° W; Fig. 1) composed of a grove of 20 cultivated *Handroanthus* spp. (Bignoniaceae) of similar ages (ca. 20 years) and growing from 2.5 to 47.5m from each other, surrounded by a grassland matrix (ca. 100 tree/ha; Fig. 1). We sampled a total of 224 individuals of *T. recurvata* growing on 14 neighboring trees (16 individuals per tree; hereafter referred to as "subpopulations") and extracted total genomic DNA from leaf samples according to Tel-Zur et al. (1999). We characterized the multi-locus genotype (MLG) of each individual using seven SSR loci designed for other species and cross-amplified them as described by Chaves et al. (2018). We avoided sampling clones by collecting individuals at distinct branches of each host tree. Indeed, clonal individuals of *T. recurvata* (i.e., ramets) can be easily distinguished because they are formed on leaf axils and remain attached, conferring the typical "ball" shape of the species (Birge 1911).

**Figure 1.**
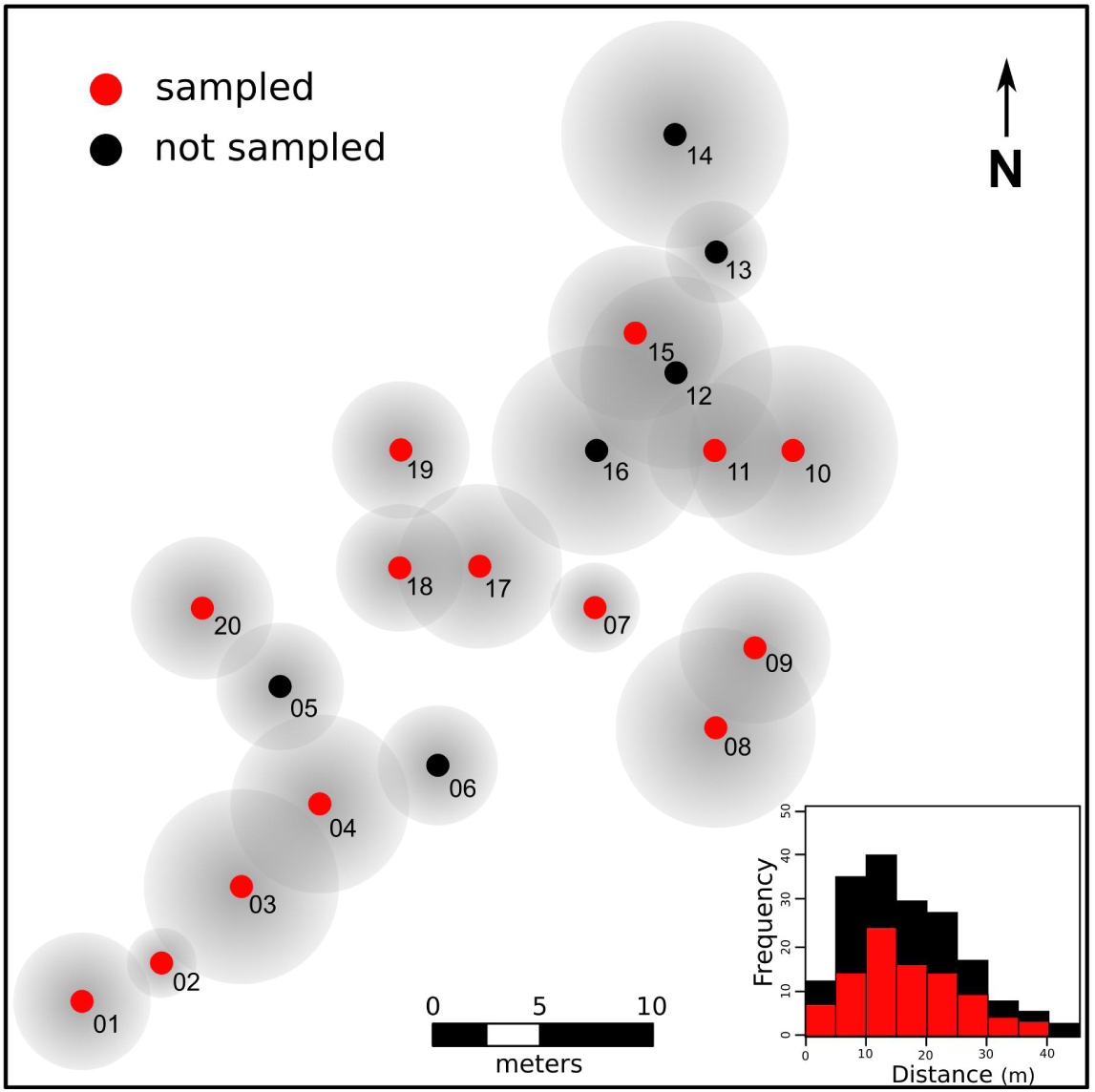
Spatial distribution and histogram of the frequency of distance classes of pairwise trees within the studied landscape. Numbers refer to IDs of trees hosting the *Tillandsia recurvata* populations analyzed. Gray circles around each dot are representations of the crown area of each tree. Red dots and bars represent, respectively, the location and frequency of distances among pairs.

To describe the genetic diversity within each *T. recurvata* subpopulation, we calculated the number of alleles (A), allelic richness (A_R_), the private number of alleles (A_P_), expected (H_E_) and observed heterozygosity (H_O_), and inbreeding coefficient (*F*_IS_) and tested for departures from Hardy-Weinberg expectations using the R statistical package diveRsity (Keenan *et al.* 2013). Furthermore, we tested if subpopulations were isolated-by-distance using a Mantel test with Slatkin's linearized *F*_ST_ (*F*_ST_ / (1 - *F*_ST_)) (1995), Nei’s (1972), Edward’s (1971), and Reynold’s (1983) pairwise genetic distances and the logarithm of geographical distance, with 1,000 replications, using the R statistical packages poppr (Kamvar *et al.* 2014) and pegas (Paradis 2010). We also measured pairwise subpopulation differentiation (*F*_ST_) and tested the hierarchical partition of genetic variance among and within sampled subpopulations by performing an analysis of molecular variance (AMOVA; Excoffier *et al.* 1992), using the R statistical package poppr (Kamvar *et al.* 2014).

To investigate the spatial genetic structure (SGS) within the *T. recurvata* population sampled, we adopted two distinct approaches, one explicitly considers the spatial distance among individuals, and another takes into account the discrete distribution of this epiphyte among distinct tree crowns. In the first approach, we tested whether the distance among *T. recurvata* individuals, disregarding tree crowns, affect their relatedness. For this, we estimated the kinship coefficient (F_ij_; Loiselle *et al.* 1995) between all pairs of individuals sampled within seven distance classes, each comprising the same number of pairs, using the R package RClone (Bailleul *et al.* 2016). To quantify the SGS under this approach, we calculated the Sp statistics (i.e. the ratio between the kinship decay by increasing geographic distances and the mean kinship within the first interval distance; Vekemans and Hardy 2004) and tested the statistical significance of mean kinship coefficient values in each distance class (confidence interval: 95%) by randomly shuffle (1000×) individual locations (Vekemans and Hardy 2004). In the second SGS approach, we rather tested whether individuals within subpopulations are more related than across the whole population. For this, we first calculated the F_ij_ for all sampled pairs of individuals within each subpopulation (F_ij_^sub^) and also for all sampled pairs in the whole population (F_ij_^pop^), using the R package RClone (Bailleul *et al.* 2016). We then calculated the ratio of F_ij_^sub^ divided by F_ij_^pop^ (F_ij_^sub^/F_ij_^pop^) for each subpopulation. Lastly, we quantified the partitioning of genetic composition of the whole population by calculating the turnover of alleles and multi-locus genotypes among subpopulations (MLG; see Harrison *et al.* 1992; Jost 2007), using the R statistical package vegetarian (Charney and Record 2012).

### Individual-based modeling

We implemented an IBM in NetLogo 6.0.1 (Wilensky 1999) and outlined it following the ODD (Overview, Design concepts, Details) protocol formulated by Grimm *et al.* (2006, 2010; see Appendix S1 in Supporting Information). The model simulates a static landscape composed of multiple 0.01 m2 patches of soil with scattered trees that are colonized by multiple seeds of *T. recurvata* with distinct MLGs (based on empirical alleles). Such seeds annually reach the landscape from outside (hereafter referred to as 'regional seeds') by wind dispersal, reproduce by self-fertilization and grow by producing new ramets (clonal growth) on trees within the landscape (Fig. 2). The wind speed was set as a modelling unit that quantifies the total distance (in units of ‘landscape patches’) seeds are carried by wind at each time step of the model. The colonization dynamics of the model is primarily based on the ‘energy budget’ of *T. recurvata*, which quantifies, in a simplified approach, the amount of energy each individual takes up by photosynthesis and expends during its life cycle through metabolism, growth, and reproduction, depending on the shading rate of its attachment site (Fig. 2). The energy budget approach is based on the Dynamic Energy Budget theory of Kooijman (2010) and has been adopted in IBMs to generally describe how individuals acquire and expend energy with simple and sufficient realism (e.g. Sibly et al. 2013; Johnston et al. 2014; van deer Vaart et al. 2016). In our study, the capacity of each *T. recurvata* individual to reproduce and generate seeds depends on the amount of energy stocked up until the reproductive season (Fig. 2). In this IBM, we assume that each seed that successfully develops and thrives in the landscape forms a multi-locus lineage (MLL) of multiple descendants, which can be traced back to the ancestor. Each new seed produced within the landscape can also be randomly wind-dispersed by chance to other trees, but new ramets are always attached to the same host tree. Each MLL often groups multiple MLGs due to heterozygous loci and mutation events. A source of genetic variation within MLLs is simulated by a varying mutation-rate (from 10^-2^ to 10^-6^) that simulates DNA replication slippages of SSR loci.

**Figure 2.**
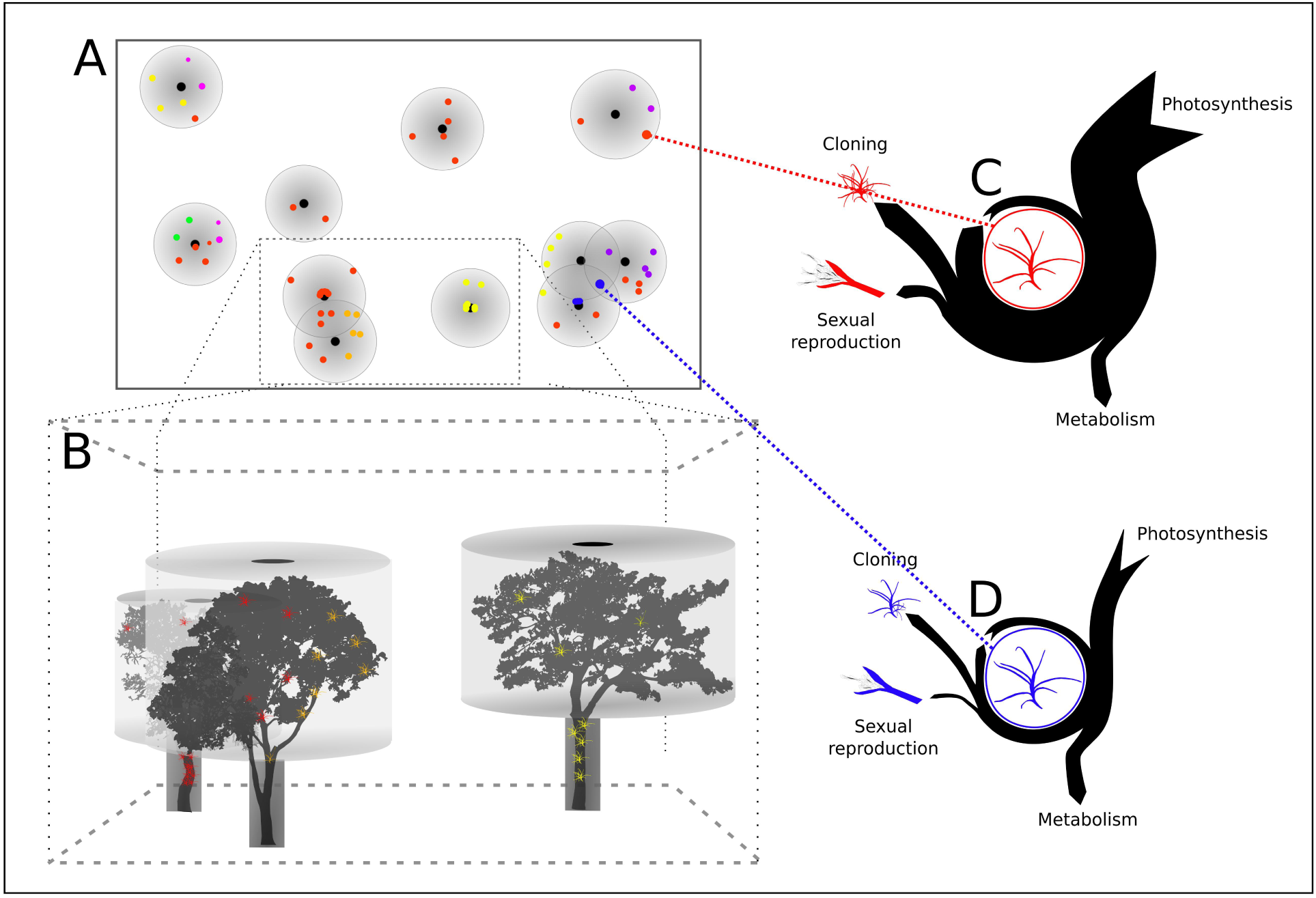
Schematic representations of the individual-based model (IBM) simulating a landscape of 0.40 hectares. Figure in A is a representation similar to what we see in the IBM in NetLogo software. Points with distinct colors represent *Tillandsia recurvat*a individuals of distinct genetic lineages (MLLs). Larger circles with a gradient gray color and concentric smaller black circles represent the crowns of distinct trees. Figure in B shows a 3D representation of the cut area of the landscape in A. Figures in C and D show the energy budget expected along a year of development of *T. recurvata* individuals attached to branches with, respectively, low and high shading levels. The arrow nocks and arrowheads represent energy inputs and outputs, respectively. The width of the arrows indicates the amount of energy being earned, lost, or accumulated.

The IBM is based on five parameters with *a priori* estimates based on general assumptions or previous knowledge on the species (see Table 1). We performed a Global Sensitivity Analysis to test whether summary statistics – i.e., the mean number of alleles (K), mean range of allele size (R), mean expected heterozygosity (H), mean inbreeding coefficient (*F*_IS_), global subpopulation differentiation (*F*_ST_), and mean modified Garza-Williamson statistic (NGW; Garza and Williamson 2001; Excoffier et al. 2005) – as well as *T. recurvata* abundance and the total number of MLL, are differently affected by the variation of each parameter. To improve the IBM model accuracy, we estimated model parameters using an ABC framework ('Approximate Bayesian Computation'; Csilléry *et al.* 2010). From a representation of the empirical landscape studied, we simulated one million SSR data sets varying five parameters, calculated six summary statistics, and obtained posterior estimates of each parameter based on 319,539 retained simulation data most resembling the empirical SSR data (see Appendix S2 for further information).

**Table 1.**
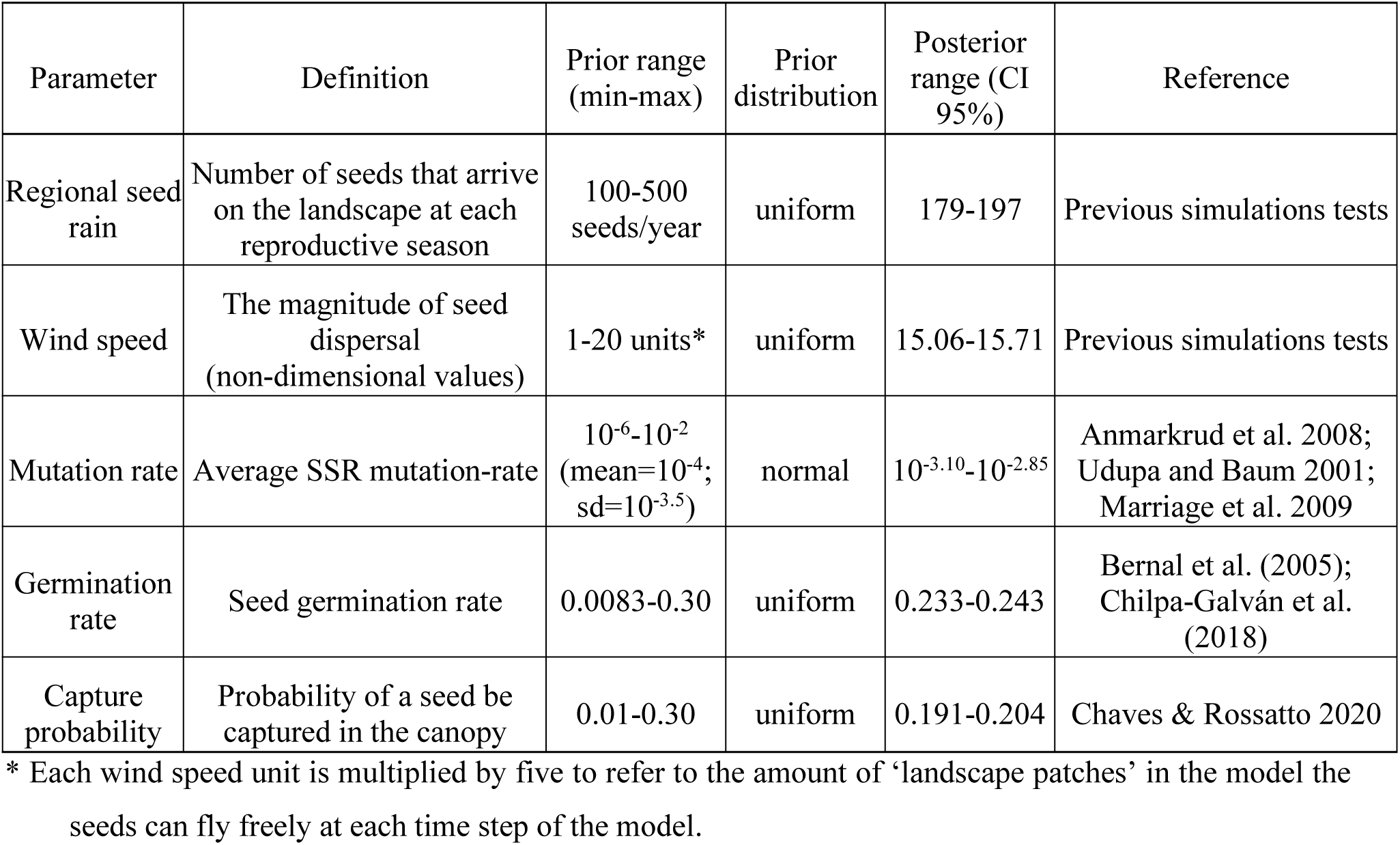
Individual-based model parameters of *Tillandsia recurvata* populations. Prior ranges are based on literature or previous simulations tests. Posterior ranges are based on the results of ABC, while estimated values are based on the accuracy of each parameter value obtained by the ABC cross-validation tests.

### Colonizing over time and across landscapes

We used the posterior range of the estimated parameters of the IBM (see Table 1) to simulate the temporal dynamic of genetic structuring during the spreading of a *T. recurvata* population (hereafter referred as ‘*time simulation*’) and the spatial dynamics of genetic structuring of populations scattered in landscapes with distinct tree densities (hereafter referred to as ‘*density simulation*’). We performed 10,000 *times simulations* for 50 years of spreading of *T. recurvata* populations on an empirical landscape representation (see Fig. 1), and 100,000 *density simulations* of *T. recurvata* populations after ten years of spreading in ca. 0.40 hectares landscapes with five to 150 trees (maximum amount of trees that occupy the landscape) with random heights and same crown sizes. After each year in *time simulations*, we recorded the individual and MLL abundances, as well as the MLG, MLL, and origins (from clonal growth or sexual reproduction; hereafter referred to as ‘*clones*’ and ‘*seeds*’ individuals) of up to 15 *T. recurvata* individuals established on each host tree representation from the empirical study. The same data were recorded after the 10^th^ year for up to 15 random trees for *density simulations*. Based on the MLGs of each simulation, we calculated the average number of alleles (A), the inbreeding coefficient (*F*_IS_), and the global subpopulation differentiation (*F*_ST_) using the ARLSUMSTAT console version of Arlequin 3.1; the MLG and MLL turnovers among subpopulations (T/O_MLG_ and T/O_MLL_), using the R package vegetarian (Harrison *et al.* 1992; Jost 2007); the kinship coefficient (F_ij_; Loiselle *et al.* 1995) of each subpopulation (F_ij_^sub^) divided by the kinship of the whole population (F_ij_^pop^) in each simulation (i.e., F_ij_^sub/pop^); and the S_p_ statistics considering seven distance classes as in the empirical study (Vekemans and Hardy 2004) using the R package RClone (Bailleul *et al.* 2016).

## RESULTS

### Empirical population genetics

We observed moderate levels of genetic diversity within each subpopulation, despite high endogamy, and high genetic structure among the studied subpopulations. The number of alleles (A) ranged from 19 to 30, while allelic richness (A_R_) ranged from 2.20 to 3.54, and the private number of alleles (A_P_) was up to 2 per subpopulation (see Table S3 in Appendix S2). Expected (H_E_) and observed (H_O_) heterozygosities ranged from 0.27 to 0.51, and from 0 to 0.20, respectively (Table S1). The inbreeding coefficients (*F*_IS_) were significant and very high, ranging from 0.52 to 1.0 per subpopulation, due to significant heterozygosity deficit under Hardy-Weinberg equilibrium (Table S1). Pairwise subpopulation differentiation (*F*_ST_) ranged from 0.01 to 0.31 (Fig. S1). The Mantel test indicated a significant although weak isolation-by-distance (IBD) pattern among subpopulations only when using Edward's genetic distances (r = 0.243; p = 0.032). AMOVA analysis, in turn, showed lower genetic variance among subpopulations (17.26%) than within subpopulations (82.74%) but was significantly greater than zero (p < 0.001).

The highest kinship (F_ij_) was observed at the smallest distance interval (up to 2.5m), which is significantly higher than expected by chance (p<0.05; Fig. 3A). Kinship coefficients peaked outside the 95% confidence interval at almost all distance classes, showing a remarkable decay at three out of four distance classes higher than ca. 13 meters (Fig. 3A), and a significant spatial genetic structure (S_p_ = 0.024; p < 0.001). The F_ij_ within subpopulations (i.e., F_ij_^sub^) was, in average, almost 15 times higher than the kinship of the population (F_ij_^pop^ = 0.002). We detected allelic and MLG turnovers of 0.031 and 0.537, respectively. MLGs were evenly spaced among subpopulations, generally with one distinct dominant MLG in most subpopulations (Fig. 3B).

**Figure 3.**
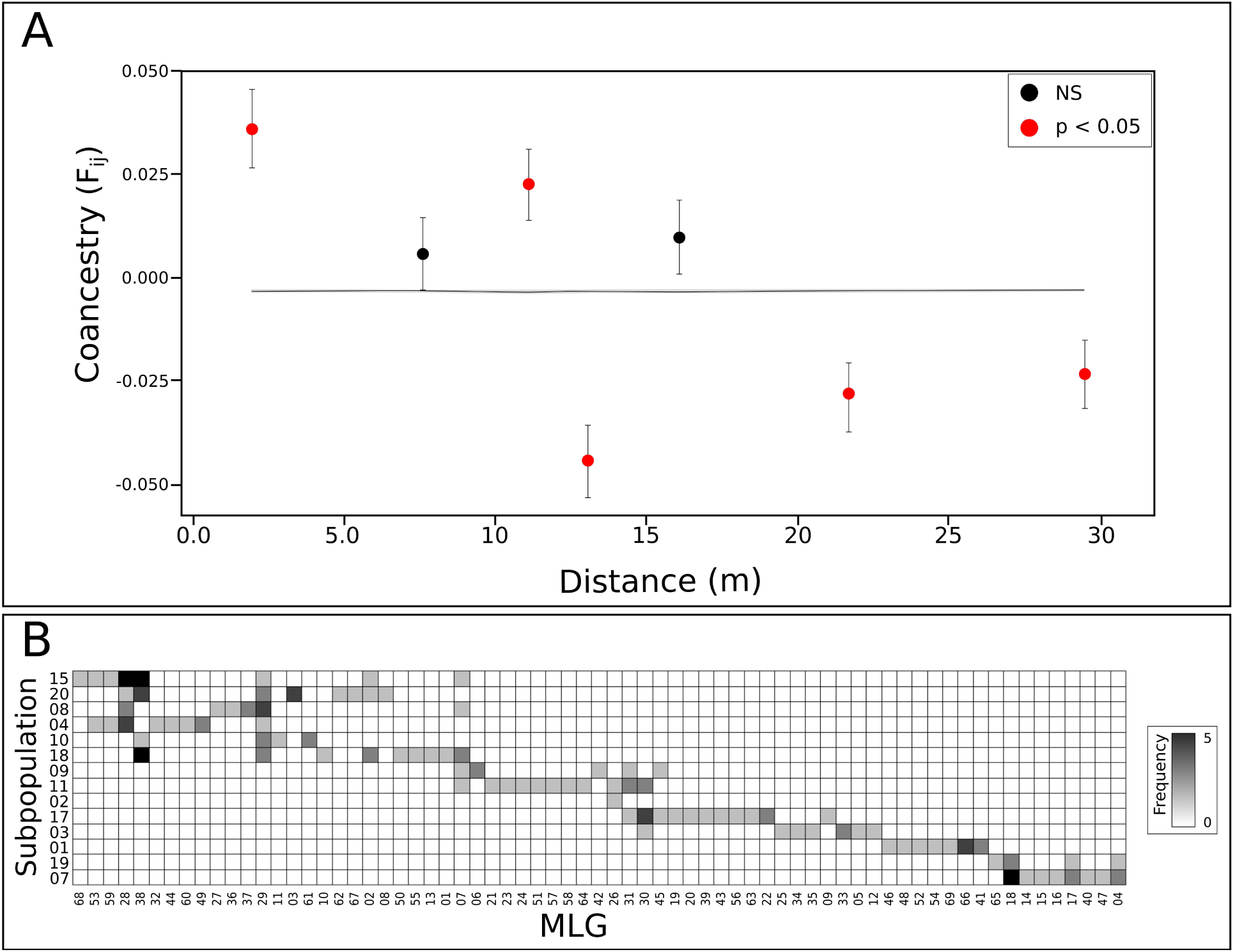
Spatial genetic structure of the empirical *Tillandsia recurvata* population. A: correlogram from spatial autocorrelation analysis using the correlation coefficient F_ij_ (Loiselle et al. 1995) and seven distance classes. The grey area represents the null hypothesis, and its 95% confidence intervals, with no spatial genetic structuring, and black lines around each average F_ij_ values represent their standard errors. B: Adjacency matrix showing the occurrence of each genotype (MLG) of *Tillandsia recurvata* (columns) on subpopulations corresponding to each sampled tree (rows). Grey gradient colors represent the frequency of each genotype per subpopulation.

### Individual-based model

The sensitivity test performed for the IBM model showed that high mutation rates of SSR markers can significantly decrease the inbreeding coefficient (*F*_IS_) and subpopulation differentiation (*F*_ST_) (Figure S2). This test also suggests that strong winds, massive regional seed rains, and high rates of germination and seed capturing likely reduce the SGS within *T. recurvata* populations (Figure S2). Based on the ABC framework, we estimated a range of parameters’ values to be randomly used in *time* and *density simulations* (Table 1). Almost all summary statistics demonstrate a deep sensitivity to parameters variation (Fig. S2). However, parameters estimation through the cross-validation showed a significant correlation between real and estimated values (p < 0.001), *wind speed* and *capture probability* were, respectively, the parameters most and least accurately predicted by the model (Fig. S3). From 319,539 simulations used for the cross-validation, this analysis accepted the parameter combination of 1,598 simulations. We found five significant correlations among parameters in accepted simulations (Table S2). *Capture probability* and *regional seed rain* (r = -0.430) are the most correlated parameters.

Individual abundance varies through time, exhibiting three distinct phases, here named Lag (ca. 0 to 5 years), Log (ca. 5 to 15 years), and Stationary (higher than 15 years) phases (Fig. 4). During the Lag phase, the individual abundance is maintained at very low values, while the MLL abundance and clonal proportion increase along with the gradual rise in *F*_IS_ and A and the decay in T/O_MLG_ (Fig. 4 and 5). Values of *F*_ST_ reach6 a maximum of ca. 0.40 in the middle of the lag phase, followed by a peak in F_ij_^sub/pop^ and Sp (ca. 20 and 0.09, respectively) and a rapid drop in F_ij_^pop^ at the end of this phase (from ca. 0.013 to 0.002; Fig. 5). Our simulations show that individual abundance multiplies, reaching a peak at the end of the Log phase (from some dozens to ca. 7,500 individuals), while S_p_ suffers large drops until the end of the stage (from ca. 0.090 to 0.045; Fig. 5). In the middle of the Log phase, the proportion of clones and *F*IS reach their maximum (ca. 0.98 and 0.80) while *F*_ST_ and T/O_MLG_ reach their minimum (ca. 0.20 and 0.50; Fig. 5). For up to 20 years of colonization (Stationary phase), simulations show that individual abundance, and many other measured variables, suffer only small variations (Fig. 4 and 5). MLL abundance and A reach to their maximum of ca. 35 MLLs and 20.5 alleles, respectively, a few years after the beginning of the Stationary phase (Fig. 5 and 6). From the Log to Stationary phases, values of F_ij_^pop^ vary between 0.0004 and 0.0013 in cycles of ca. of 4 years.

**Figure 4.**
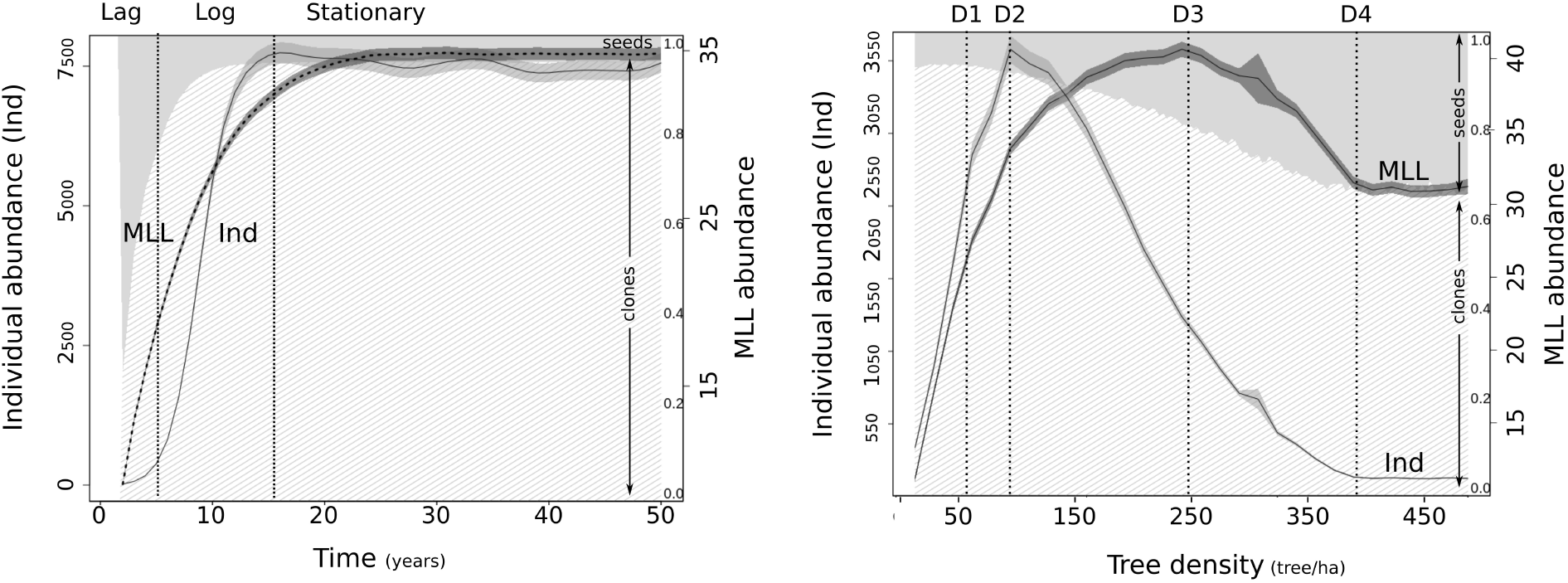
Variation of individual and multi-locus lineages (MLL) abundances of *Tillandsia recurvata*, as well as its clones/seeds proportion, according to time of colonization and tree density. Stages of individual abundance development by time are highlighted as “Lag”, “Log”, and “Stationary”. Critical tree densities for parameters variation are highlighted as “D1”, “D2”, “D3”, and “D4”.

**Figure 5.**
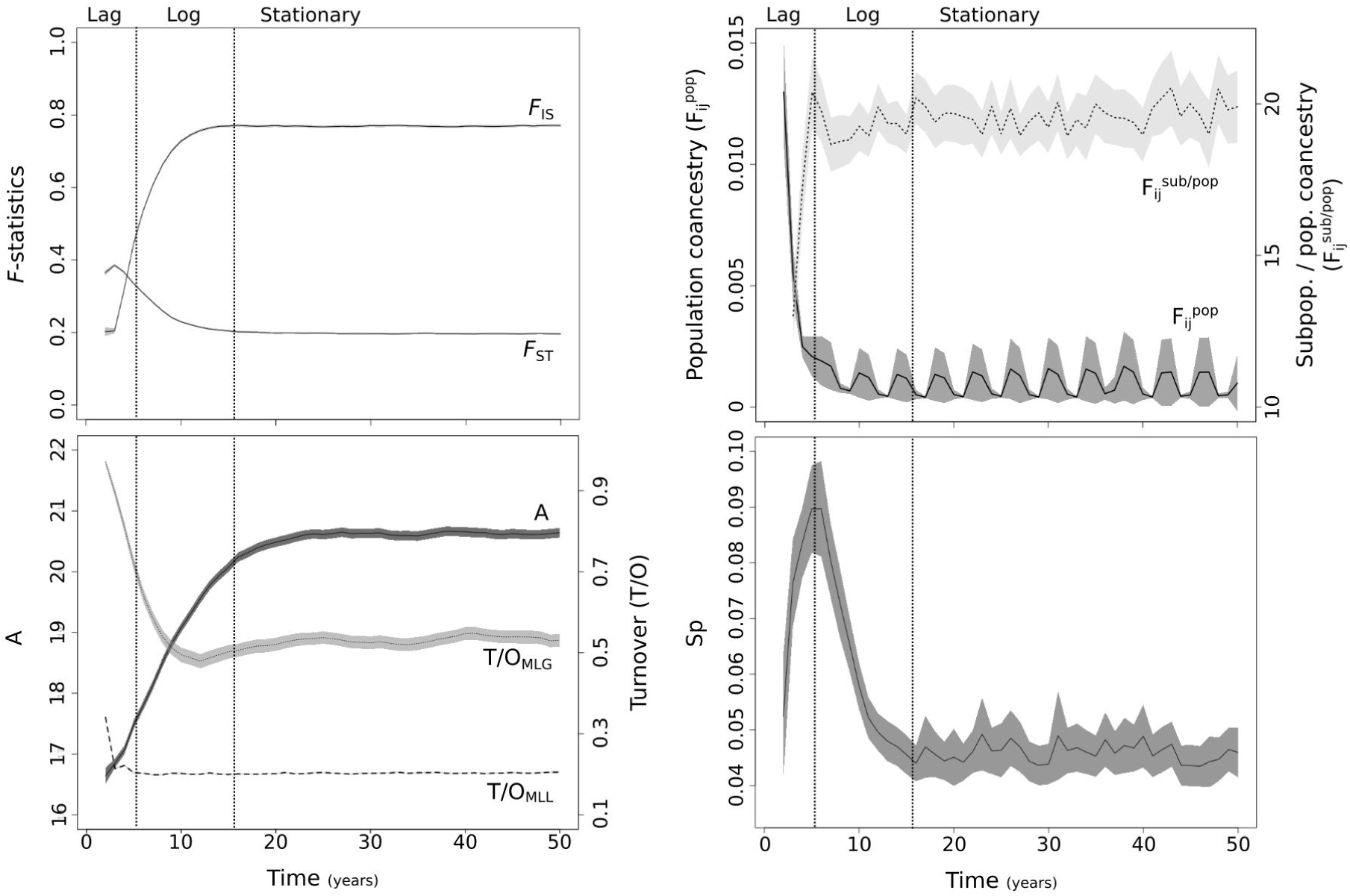
Temporal dynamic of genetic indices of *Tillandsia recurvata* population. Lines and grey areas around them represent the average and 95 % confidence intervals of 10,000 simulations. Stages of individual abundance development by time are highlighted as “Lag”, “Log”, and “Stationary”.

**Figure 6.**
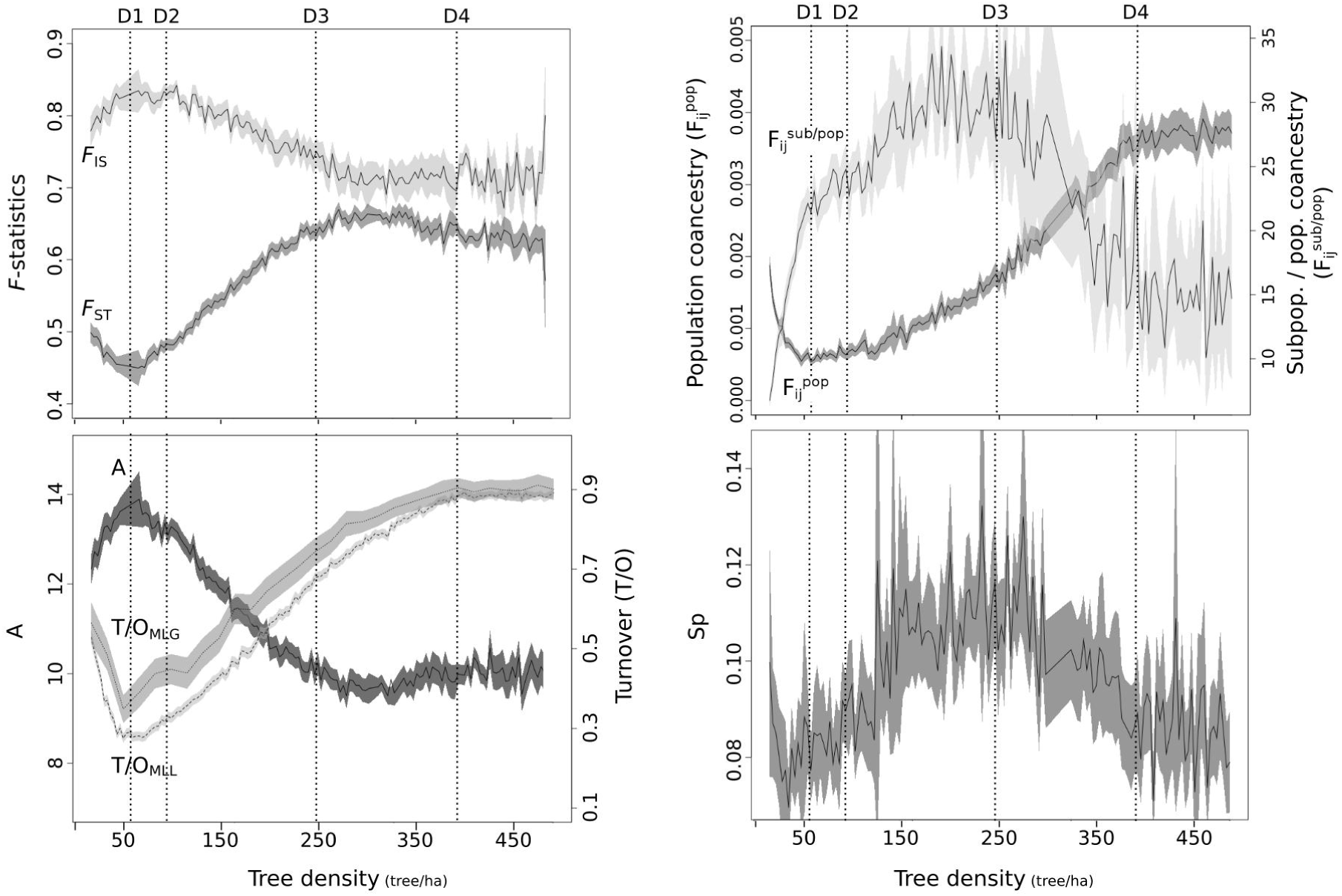
The spatial dynamic of genetic indices of *Tillandsia recurvata* population over landscapes with distinct tree densities. Lines and grey areas around them represent the average and 95 % confidence intervals of 100,000 simulations. Vertical dotted lines highlight key values (D1-D4) of tree densities according to the studied indices.

*Density simulations* show greater variance and less stability than *time simulations*. We found that increasing tree densities of up to ca. 50 trees/ha (D1 in Figs. 4 and 6) lead to the greatest proportion of clonal individuals (ca. 0.90, see Fig. 4) and a peak of *F*_IS_ (ca. 0.85) and the number of alleles (ca. 14). At the same time, *F*_ST_, T/O_MLG_, T/O_MLL_, F_ij_^pop^, as well as the Sp statistics reached their lowest values (ca. 0.45, 0.35, 0.28, 0.0005, and 0.075 respectively, see Fig. 6). Our simulations also showed that the peak of individual abundance (ca. 3600 individuals) occurs at ca. 95 trees/ha (D2 in Fig. 4) and that increasing tree density up to 400 trees/ha (D4 in Fig. 4) leads to a reduction of up to 95% of individuals. Intermediate tree densities of ca. 250 trees/ha (D3 in Fig. 4) foster the greatest MLL abundances (ca. 40 MLLs), and the highest values of *F*_ST_ (ca. 0.65), F_ij_^sub/pop^ (ca. 30), and Sp statistics (ca. 0.11). A steep decay in the proportion of clones (from ca. 0.95 to 0.65), between 50 and 400 trees/ha (D1 and D4 in Fig. 4), is followed by the decay of *F*_IS_ (ca. 0.85 to 0.70) and number of alleles (ca. 14 to 10), and the increase of *F*_ST_ (ca. 0.45 to 0.65), T/O_MLG_ (ca. 0.35 to 0.90), T/O_MLL_ (ca. 0.25 to 0.90), and F_ij_^pop^ (ca. 0.0005 to 0.0040).

## DISCUSSION

In this study, we integrated empirical and simulated genetic data to provide insights into the dynamics of genetic structuring of populations of an epiphytic weed (i.e., *Tillandsia recurvata*) distributed into human-transformed landscapes. Our empirical results show a relatively high spatial genetic structure (SGS) for *T. recurvata* within a small anthropic landscape (c.d. 0.2 ha) with low tree density, suggesting that trees may be considered as habitat units of epiphytic systems (Flores-Palacios & García-Franco 2006). The simulated data provided by our individual-based model (IBM) explicitly describes the temporal dynamics of the empirical *T. recurvata* population, revealing the highest levels of SGS at the very beginning of the colonization process. The IBM also points out that the landscapes with the highest tree densities tend to harbor populations of *T. recurvata* with the strongest SGS patterns. Below, we discuss our findings based on the ecological features of this species, highlighting the importance of understanding the spatio-temporal dynamics of genetic structuring and the eco-evolutive consequences of human transformations on landscapes.

### Fine-scale genetic structure

Our empirical study in an anthropic landscape shows a relatively pronounced SGS for *T. recurvata* at a fine-scale (ca. 2,000m^2^), as indicated by the significant S_p_ statistics, high subpopulation differentiation, and the almost 15× greater kinship among individuals within subpopulations compared to the kinship in the whole population. Such SGS likely arises as an outcome of the breeding system and life form of *T. recurvata*, which present limited local gene flow, resulting in a significant isolation-by-distance, persistence of private alleles in six out of 14 studied subpopulations, as well as high turnover of multi-locus genotypes (MLGs). Indeed, SGS at fine scales often results from limited gene flow (Vekemans and Hardy 2004; Ward 2006), and other studies have reported similar genetic patterns for selfing (e.g., Pettengill *et al.* 2016; Atwater *et al.* 2018), cloning (e.g., Li and Dong 2009; Atwater *et al.* 2018), epiphytic (e.g., Trapnell *et al.* 2004; Ren *et al.* 2017; Torres *et al.* 2018), and fast-growing plants (e.g., Barluenga *et al.* 2011; Charbonneau *et al.* 2018). Our results indicate that specific habits and breeding systems can foster the structuring of populations, even for opportunistic and widespread plants.

Our results also indicate that the *T. recurvata* spreading is contemporary and gradually conducted by several sources following a phalanx pattern, rather than by a single leading subpopulation (see Hewitt 1996), considering that many MLGs have the same alleles. Still, few subpopulations share the same MLGs (i.e., low allelic turnover but high MLG turnover). These results may suggest that distinct native populations of *T. recurvata* of the surroundings may be responsible for colonizing our focal landscape massively. Under such conditions, selfing reproduction and clonal growth may lead to a partial, and perhaps transitory, isolation among subpopulations of *T. recuvata* (see Ward 2006). Beyond the effect of breeding system and life form, the relatively high fine-scale SGS of *T. recurvata* may also result from following founder events on each host tree (see Hewitt 1996; Ward 2006). Nevertheless, the genetic drift resulting from consecutive founder events, as well as the increasing amount of seeds released from large subpopulations, may intensify the metapopulation dynamics of repeated local extinction and recolonization events, which may reduce the population structuring over time (Baker 1955, Pannell and Barrett 1998; Razanajatovo *et al.* 2016).

### SGS over time

The genetic structure dynamics of *T. recurvata* throughout the *time simulations* of our IBM, as represented by the Lag, Log, and Stationary phases for distinct genetic parameters, resembles diffusion models of invasion for weedy species in new disturbing landscapes (e.g., Counsens and Mortimer 1995). In the first few years (i.e., Lag or stationary phase; here, up to the ^5th^ year), only a few individuals successfully establish in the landscape, despite the constant input of new genetic lineages (MLLs) per reproductive season. The mismatch between the small number of *T. recurvata* individuals and the increasing number of MLLs reduces the average kinship in the population and leads to a peak of genetic structuring and differentiation among subpopulations. The continuous establishment of new MLLs within the landscape underlies an increasing production of new ramets by clonal growth, that may help to overcome demographic stochasticity and the Allee effect (Loreau at al. 2013; Santos *et al.* 2014; van Kleunen 2018), raising the individual abundance and inbreeding coefficient (*F*_IS_) to their highest values in the Log phase (between the 5^th^ and 15^th^ year).

The rapid increase in individual abundance during the log phase drives the interchange of seeds and MLGs among subpopulations, leading to a reduction in subpopulation differentiation and structuring. However, the stabilization in the proportion of clones in this phase limits the establishment of a higher number of individuals and MLLs, leading to a steady level in all calculated variables by the 15^th^ to 20^th^ year (Stationary phase). According to our results, the reduction in subpopulation differentiation and structuring during the Log phase, followed by stabilization at lower values, may indicate that *T. recurvata* meta-populations behave as a source population after the 15^th^ year of colonization, supporting their rapid spreading into potential new areas. Notwithstanding the specific characteristics of each species, this temporal pattern is typical for other opportunistic organisms under optimal conditions. As observed for *T. recurvata*, a fast increase and stabilization in the individual abundance of populations and communities in static systems are expected (see Spruch *et al.* 2019). On the other hand, a landscape with growing trees may lead to a gradual increase in epiphyte abundance with no signs of saturation (Einzmann and Zotz 2017; Spruch *et al.* 2019). But, given the preference of *T. recurvata* for dry and open environments, canopies with increased shading, as provided by dense forests or a reforestation system, could have an opposite effect, as we show the following.

### SGS across landscapes with distinct tree densities

Results from simulations under distinct tree densities show that the highest abundance of *T. recurvata* individuals occur at tree densities close to those observed from the empirical data (ca. 100 trees/ha). Lower tree densities (<100 trees/ha) led to a decrease in SGS may due to a reduction in habitat amount, since each host tree counts as a habitat patch for the species (e.g., Snäll *et al.* 2005; Belinchón *et al.* 2017). On the other hand, under higher tree densities (>50 trees/ha), the increasing number of trees may maximize the SGS in *T. recurvata* due to a strengthening of the resistance for seed dispersal among subpopulations. Under these conditions, winds are typically stronger and support higher wind-dispersal distances in open areas (see Nathan *et al.* 2002; Cousens *et al.* 2008). Indeed, studies implementing IBM approaches for distinct biological systems have shown that habitat amount and resistance to gene flow across landscapes strongly influence SGS among subpopulations (e.g., Cushman *et al.* 2012; Jackson and Fahrig 2016). However, even under higher tree densities (>250 trees/ha), *T. recurvata* may have reduced levels of genetic structuring (S_P_) and kinship differentiation among subpopulations (F_ij_^sub/pop^) likely due to a drop in the overall individual abundance and an underlying increase in the average kinship in population. Moreover, our simulations show that very few genetic lineages of *T. recurvata* successfully establish only at the edges of the landscapes with the highest tree densities (~450 trees/ha). Consequently, the established subpopulations often have individuals of the same lineage and, therefore, low genetic differentiation among each other.

The stronger spatial genetic structure of *T. recurvata* populations in landscapes with high tree densities is likely a consequence of the species preference for dry and open ecosystems (e.g., Smith *et al.* 1989; Orozco-Ibarrola *et al.* 2017; Chilpa-Galván *et al.* 2018). Indeed, *T. recurvata* populations are often reduced to a few individuals constrained to drier and upper layers of dense forests (Vergara-Torres *et al.*, 2010; Ruiz-Cordova et al. 2014; Flores-Palacios *et al.* 2015a; Flores-Palacios et al. 2015b). In landscapes with high tree densities, as in dense forests or reforestation systems, the high genetic drift promoted by a stronger resistance for wind dispersal (Barrett & Kohn 1991), in addition to the reduced fitness of *T. recurvata*, led to small, transient and isolated populations with low genetic diversities. The drastic reductions in tree densities induced by human-modifications in natural landscapes may have led to a partial loss of resistance for seed dispersal by wind and increased the conditions for the optimal development of *T. recurvata*. This dynamic may explain the massive abundance of *T. recurvata* individuals at lower tree densities. The high proliferation of *T. recurvata* at low tree densities suggests that human-induced deforestation plays an essential role in driving the rapid spread of opportunistic plants within landscapes that once limited their proliferation and gene dispersal. As pointed out by our sensitivity tests, the combined effects of massive regional seed rains and high rates of germination and seed capturing may explain the successful invasion of *T. recurvata* in human-transformed landscapes, besides its transitory and relatively high genetic structure (e.g., Caldiz & Fernández, 1995; Flores-Palacios *et al.*, 2014).

### Conclusion

Opportunistic species provide an excellent opportunity to examine the process of successful colonization in novel environments (van Etten at al. 2017). Our results showed that the effects of high spatial genetic structure, reduced local genetic diversity, and inbreeding, due to selfing and clonal growth of weeds (e.g., Heywood 1991; Barrett 2015; Razanajatovo *et al.* 2016), can be overwhelmed by a rapid spread across optimal open landscapes. We also demonstrate here that the reduced tree densities, commonly found in human-transformed landscapes, may have created suitable conditions for the rapid spread of *T. recurvata* into areas that might be previously limiting seed dispersal and establishment. Therefore, the reforestation of landscapes dominated by weeds with similar requirements could be an effective strategy to control their spreading.

## ACKNOWLEDGMENTS

This research is part of the first author’s thesis (Chaves 2019) and was supported by resources supplied by the Center for Scientific Computing (NCC/GridUNESP) from São Paulo State University (UNESP). We thank A. Santos for computing support. We thank S. Kakazu and M. Bacc for genotyping procedures. C.J.N.C received Ph.D. scholarships from CAPES and FAPESP (2016/04396-4; BEPE 2017/01559-2); B.S.S.L was funded by FAPESP (2014/08087-0); D.R.R. and C.P.S. were funded by Conselho Nacional de Desenvolvimento Científico e Tecnológico (CNPq 471756/ 2013-0 and 300819/2016-1, respectively).

## DATA AVAILABILITY

The microsatellite data generated in this study, as well as all scripts used in the simulations, are stored at FigShare repository [doi:10.6084/m9.figshare.12797846].

## AUTHOR CONTRIBUTIONS

CJNC conceived and designed the research, designed the IBM, proceeded and analyzed the empirical and simulated data, and wrote the draft manuscript. BSSL contributed to the simulated analysis and discussion. BSSL and CPS contributed to empirical study and discussion. DRR contributed to the empirical material collection and design. UB contributed to the model conception. All authors contributed to manuscript revision, read, and approved the submitted version.

