## Appendix S1 for "Deforestation reduces the genetic structure of an epiphytic weed across space and time: an IBM approach"

**Table S1.** Sample number (N), number of alleles (A), allelic richness (A_R_), private alleles (A_P_), expected heterozygosity (H_E_), observed heterozygosity (H_O_), and fixation index (F_IS_) for each sampled population.

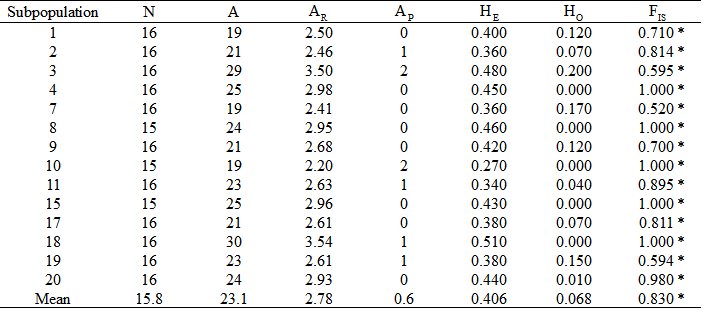

* Deviations from Hardy-Weinberg Equilibrium (P<0.05).

**Table S2.** Correlation among IBM parameters for the retained simulations. Correlation coefficients are shown in the upper triangle and their p-value are shown in the lower triangle of the matrix.

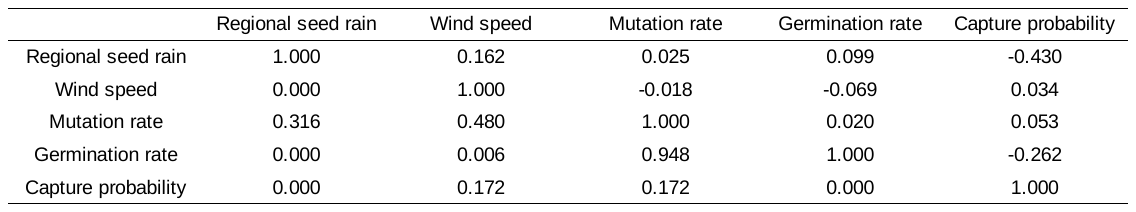

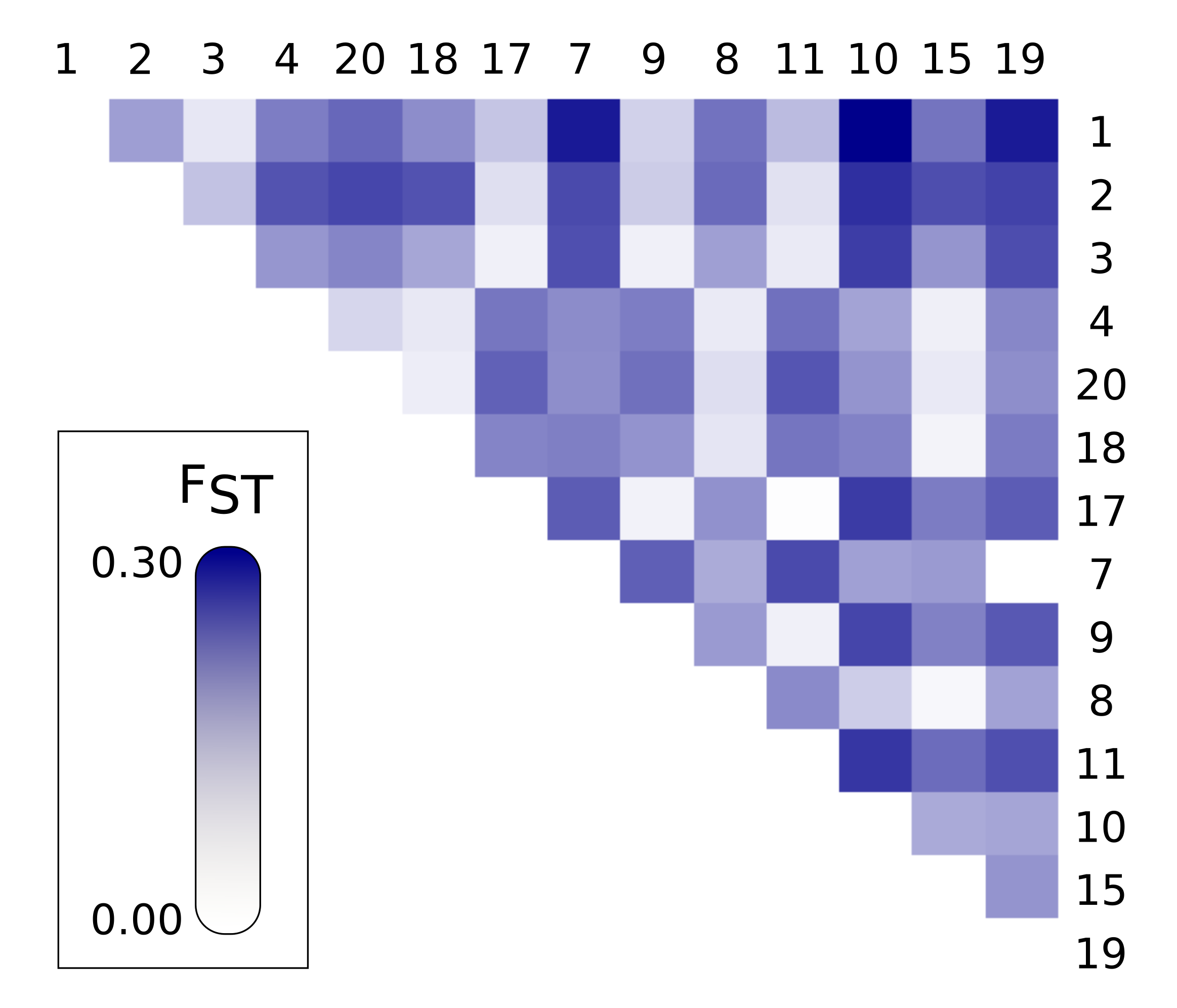

**Figure S1.** Squared matrix representation of subpopulation differentiation of *Tillandsia recurvata.*

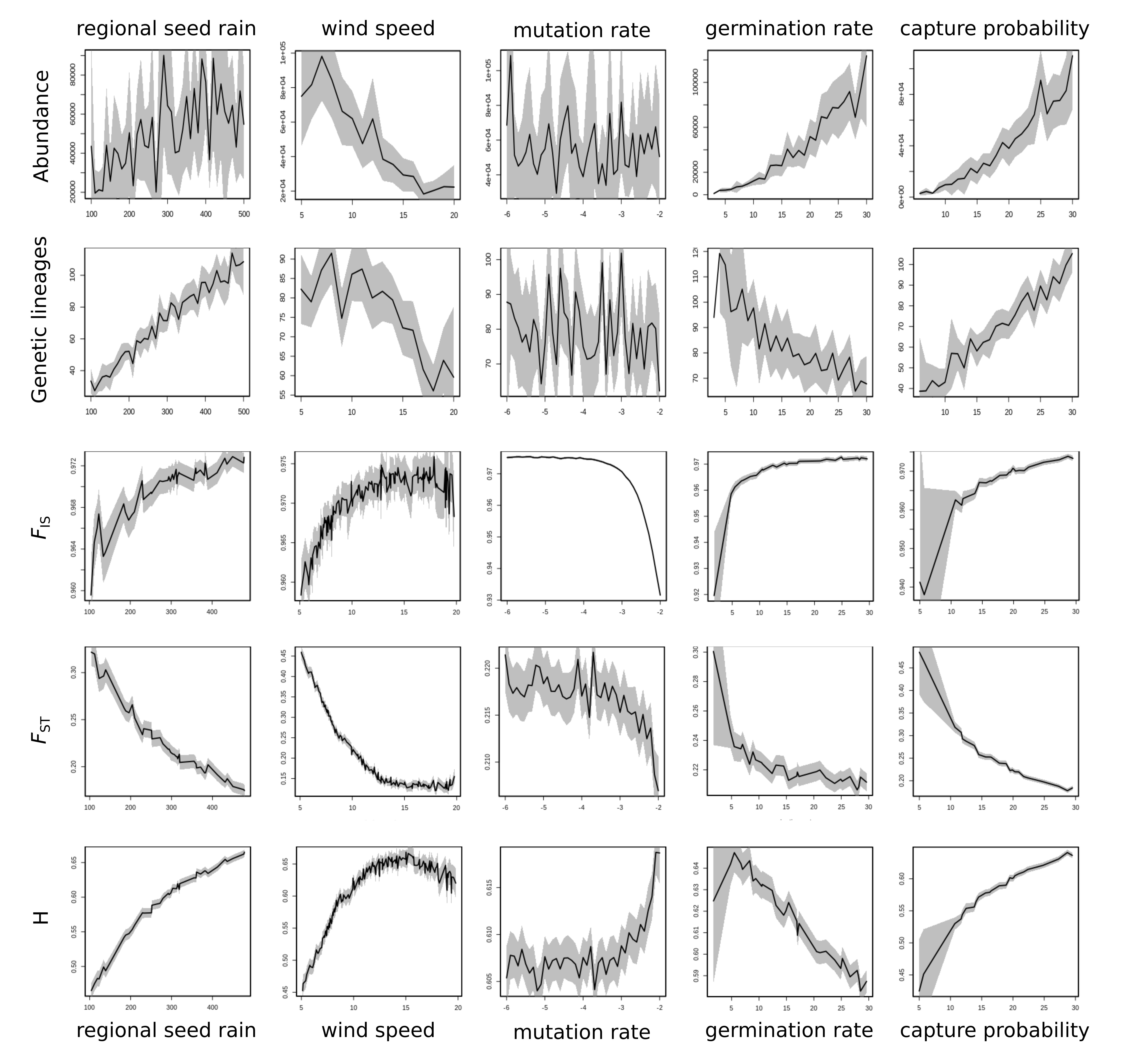
**Figure S2**. Sensitivity analysis of the IBM model. X-axes represent the parameters’ variation; Y-axes represent the summary statistics, as well as *T. recurvata* abundance and total number of genetic lineages. F_IS_: mean fixation index; F_ST_: mean population differentiation; He: mean expected heterozygosity.

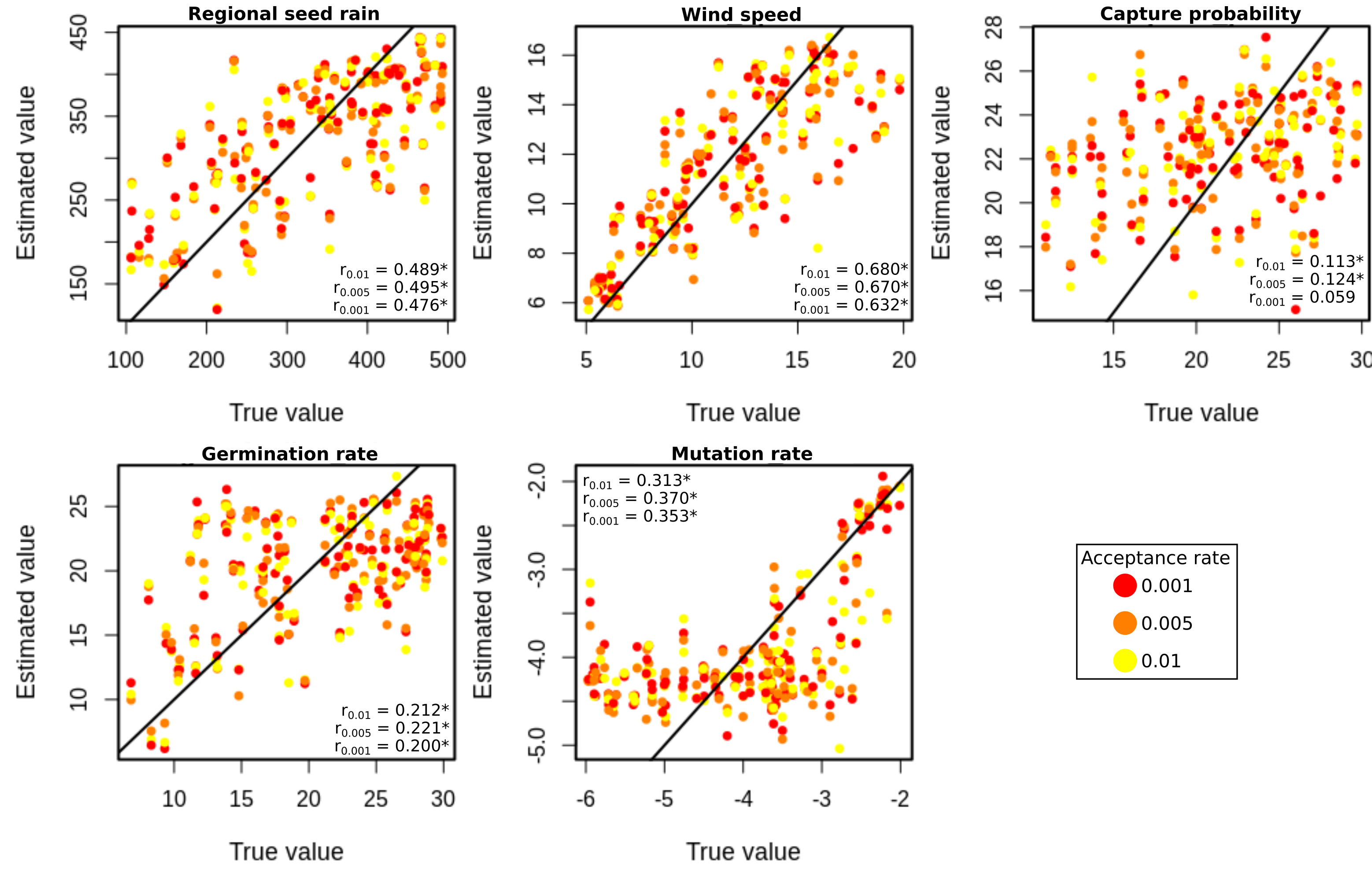
**Figure S3.** Cross-validation for parameter estimation. Parameter values estimated by ABC in relation to true values for each of the five IBM parameters. Spearman’s correlation coefficients (r) are shown at each panel for each acceptance rate.
