## Appendix S2 for "Deforestation reduces the genetic structure of an epiphytic weed across space and time: an IBM approach"

***ODD (overview, design concepts, details) of the IBM model for the establishment of a Tillandsia recurvata (L.) L. populations***

1. *Purpose*

This model simulates the colonization of a group of trees by seeds of one of the most abundant and distributed atmospheric bromeliad species, *Tillandsia recurvata* (L.) L., with distinct multilocus genotypes (MLG). This epiphytic bromeliad occurs in the American continent, from Argentina to the south of U.S.A. (Smith and Downs 1977; GBIF 2017), and is able to form populations of hundreds of individuals in trees and shrubs of native and anthropic environments (e.g. Birge 1911; McWilliams 1992; Flores-Palacios et al., 2015). Despite its huge distribution, individuals of *T. recurvata* can produce just a few dozens of seeds in each reproductive season and many of them are sterile. On the other hand, the species has a rapid life cycle, cleistogamous flowers with autonomous self-fertilization, wind-dispersed seeds, and the ability for intense clonal reproduction to group dozens of ramets that stay linked together in a ball-shaped genet producing their own seeds in each reproductive season (Soltis et al. 1987; Smith et al. 1989; Orozco-Ibarrola et al. 2015; Chilpa-Galván et al. 2018).

With this model, we propose to understand the emergence of the observed patterns obtained by our empirical study with 14 sampled trees from a grove of 20 individuals of *Handroanthus* spp. (Bignoniaceae) of similar ages (ca. 20 years) and growing from 2.5 to 47.5m from each other, surrounded by a grassland matrix (ca. 100 tree/ha; Fig. 1). We aimed on transcend the emerged pattern by trace the past and forecast the future of the empirical *T. recurvata* population and simulate landscapes with distinct tree densities. It is hypothesized that the observed spatial genetic structure (SGS) in the real landscape is due to the higher probability of seeds to be attached on the same tree where they were produced, compared to the probability to attach to other trees, forming populations with low genetic diversity. However, these results could have distinct outcomes throughout colonization time and in landscapes with different tree densities (Fig. 1). At the very beginning of *T. recurvata* spreading over a new landscape, SGS should be the strongest, but it reduces according to the offspring of the first arriving seeds spread over the trees and become a single large population (Fig. 1a). SGS could also change under distinct tree densities. The overlap of tree crowns in areas with intermediate tree densities, for instance, could acts as a bridge for the seed capturing, but the massive overlap among these crowns, under high tree densities, could acts inversely, reducing the local wind speed and shading, which hamper seed dispersal, increase the boundary layer and reduce
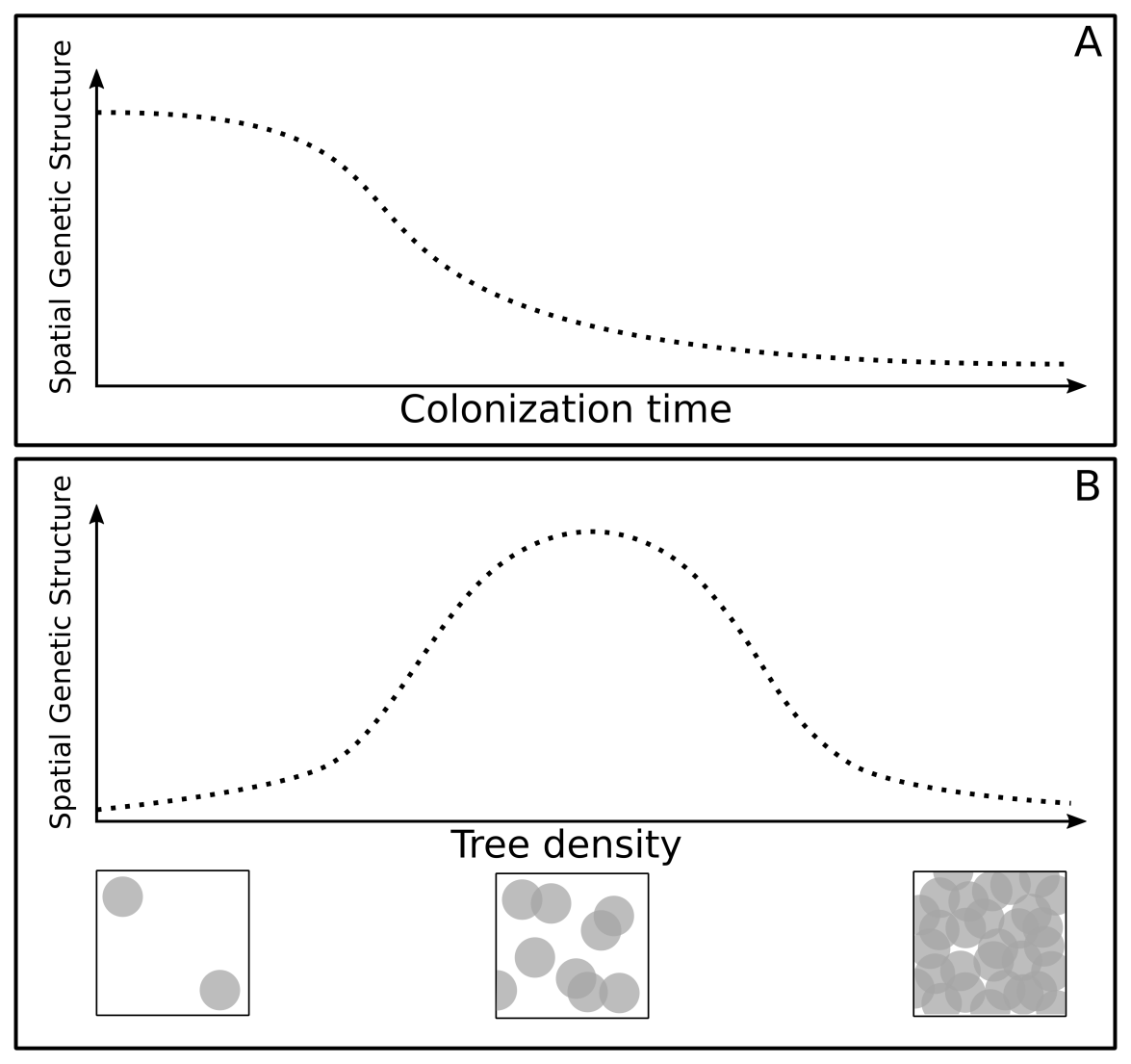
photosynthesis rate (Fig. S1b). Since *T. recurvata* is a drought adapted species (e.g. Benzing 2000; Benzing 2012), these environmental conditions could lead to a reduction in the species’ population and increase its mortality. Conversely, the seed capturing in areas with very low tree densities could also be hampered, since there is less substrate for the attachment of seeds, also increasing SGS (Fig. 1b). At the same time, all this SGS dynamics will also depend on the mutation rate of *T. recurvata* markers and on: (i) the wind power to carry the seeds of *T. recurvata*; (ii) the amount of seeds from the regional seed rain that reaches the landscape at each year; (iii) the germination rate of *T. recurvata* seeds; and (iv) the probability of seeds captured by potential host trees.

Figure 1. Schematic representation of the relationship of spatial genetic structure of Tillandsia recurvata population with the colonization time of a landscape (A), and with the tree density of landscapes. The squares in B represent landscapes with distinct numbers of trees (gray circles) at a top view looking down.

2. *Entities, state variables, and scales*

This model follows the growth and spreading of a *T. recurvata* population on trees scattered in a two-dimensional landscape at monthly intervals. The landscape area is comprised by multiple
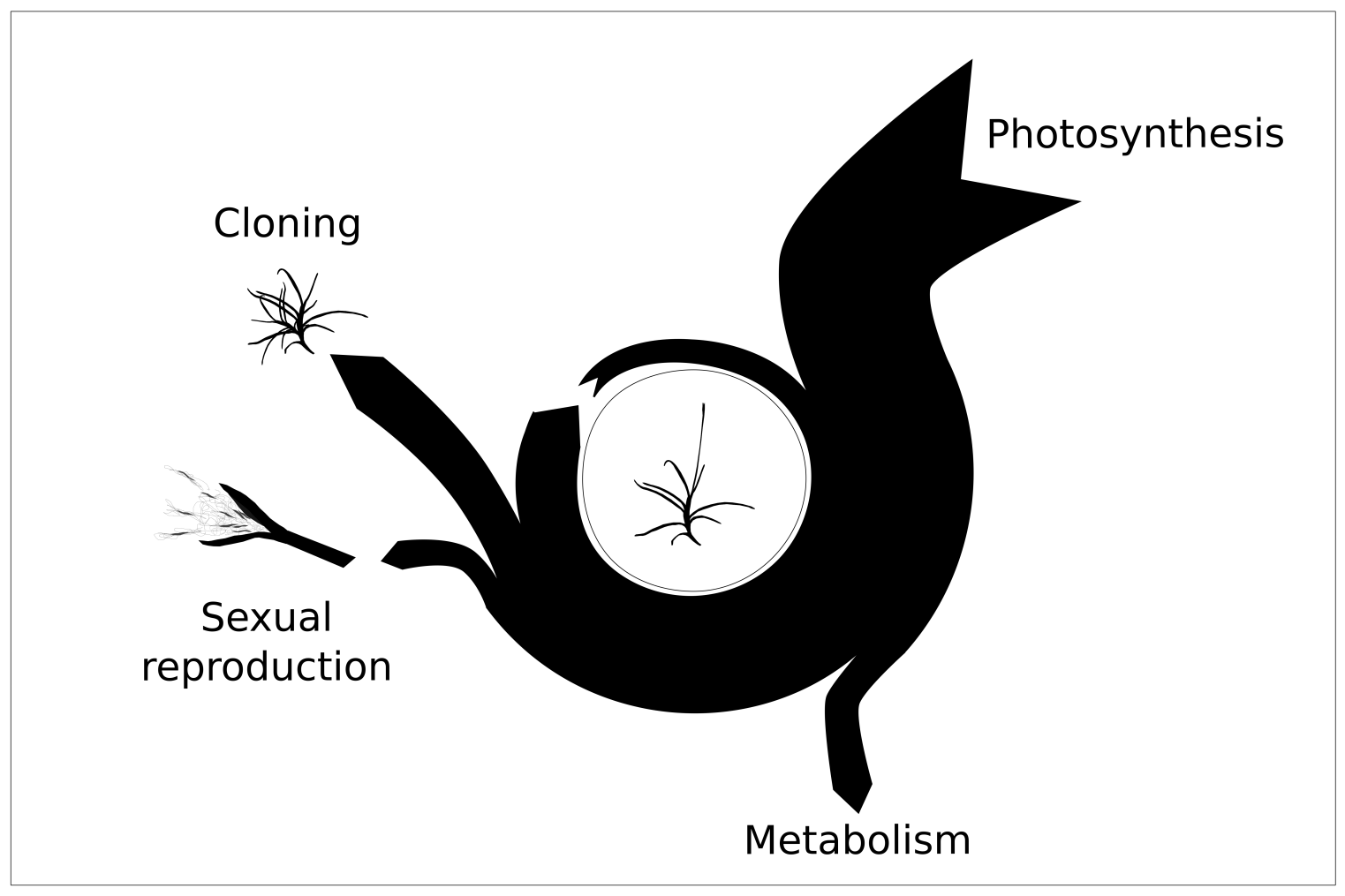
0.01 m^2^ patches of soil, and can be a representation of both an empirical or artificial landscape. The empirical landscape has trees with predefined sizes and positions (Table 1) scattered in the ca. 2,000 m^2^ area where we performed the empirical study with *T. recurvata* populations. The artificial landscape has a squared area of ca. 0.40 ha. with randomly distributed trees with previously specified sizes. Each tree in the landscape has its own size (in meters), which is divided in trunk height, crown height, and crown area. Individuals of *T. recurvata* are characterized by their age (in months), lifespan (in months), height of attaching on the host tree (in meters), origin (“seed” or “clonal ramet”), location on the host tree (“canopy” or “trunk”), genotype (comprising two alleles of each of the seven SSR markers used in the empirical study), and energy (in arbitrary units). To simulate the effect of shading levels and competition on the fitness of each atmospheric bromeliad, we simulated their energy budget that calculates their energy gains by photosynthesis and their expenses by metabolism and reproduction (Fig. 2).

**Figure 2.** Representation of the simplified energy budget of an individual of *T. recurvata*, in which all model is based. The arrow nocks and arrowheads represent, respectively, energy inputs and outputs during a cycle. The width of the arrow indicate the amount of energy being earned, lost, or accumulated and is variable according the local shading rate.

**Table 1**. Coordinates and features measured on the trees of empirical landscape.

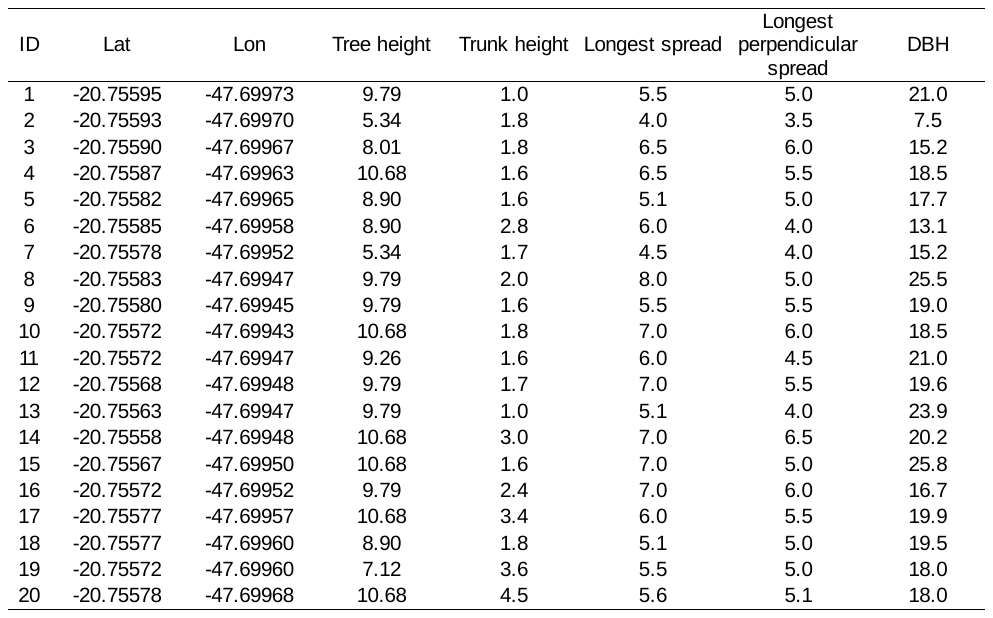

3. *Process overview and scheduling*

Based on scientific literature and personal observations on the life cycle of *T. recurvata*, the following list of processes is executed once per time step (i.e. monthly):

1. **Update:** The model updates the age in months of all *T. recurvata* individuals and calculates their energy budget (Fig. S2). The latter is based on the gains and expenses by photosynthesis and metabolism taking into account the shading rate, local space sharing, and individual age.
2. **Reproductive season:** Every year (i.e. corresponding to 12 time steps) the model adds new bromeliad seeds to the system. These seeds come from outside of the system (hereafter referred as ‘regional seed rain’) or from reproducing individuals already attached to trees of the system (hereafter referred as ‘local seed rain’). The regional seed rain is initially the main source of genetic diversity for subpopulations (i.e. individuals attached to the same tree), as each new introduced seed has a random genotype where the alleles and heterozygosity of each locus follows the observed in the empirical data. The genotypes of seeds from local seed rain, in turn, are a combination of the maternal gametes’ haplotypes, which contributes to increase the abundance of local multilocus genetic lineages (MLLs). Here, MLLs are referred as a group of individuals in which the common ancestor is the first seed that arrived from the regional seed rain. Nevertheless, locally produced seeds can also increase the genetic diversity of a population, since they are subject to mutations by DNA replication slippage. Moreover, adult individuals could also reproduce vegetatively, producing new ramets that stay attached to them and have the same genotypes. To reproduce, vegetatively or through seed dispersal, bromeliads have to have enough energy at the reproductive season. This energy is acquired along the year and is balanced with the energy spent on metabolism. For this reason, it is reduced in stablished bromeliads under heavy competition or strong tree crown overlapping (i.e. shading rate).
3. **Seed movement:** Dispersed seeds move straightforward in a random direction, gradually reducing their height. They can be captured if they are on the same height and location of a tree trunk or crown. If, by chance, the seed is not captured by the canopy, its velocity reduces drastically, due to the weaker wind speed inside canopies (see Nathan et al. 2002). Seeds that reach the ground (height = 0) or to the margins of the simulated landscape are excluded from the model.
4. **Seed germination:** The model calculates the germination probability of seeds in the month after been being captured by a tree trunk or crown. If not germinated, they are excluded from the model.
5. **Mortality:** If a bromeliad reaches its lifespan age or if accumulated energy reaches to zero, it is also excluded from the model.

*4. Design Concepts*

- *Basic principles*

We assume that *T*. *recurvata* is a drought-tolerant epiphytic species with a cleistogamous reproductive system, which has closed flowers that self-fertilize to produce new seeds. Under this view, *T. recurvata* has ‘closed MLLs’, where the genetic diversity in a population increases only by the arrival of seeds from a new lineage or by the mutation of locally produced offspring.

- *Emergence*

The genetic structure of the simulated *T. recurvata* population emerges from the reproductive and seed dispersal processes, and from the parameters set on the setup procedure (i.e. wind speed, regional seed rain, seed capture probability, mutation rate, and tree density). It is measured with *a posteriori* analysis that uses the genotypes of up to 15 randomly sampled bromeliads composing each of the 14 subpopulation sampled in the empirical landscape or up to 15 subpopulations formed in simulated landscapes.

- *Stochasticity*

If the landscape is set as artificial, the position of trees is chosen randomly from the unoccupied patches. Regional seeds enter in the landscape from a random point in its margins, or from points of the corresponding margin from which the direction of regional seed rain is set. If activated, disturbance events occur randomly once each five months, harming all bromeliads and reducing drastically their stored energy.

- *Observation*

A 2D-display produced in NetLogo software (version 6.1; Willensky, 1999) allows for the monitoring of *T. recurvata* population spreading in the simulated landscape. It shows a top view looking down on a landscape formed by trees, represented by green circular crowns with a trunk shown as a brown and smaller concentric circle. Moving colorful points represent dispersing *T. recurvata* seeds which develop into seedlings and adults when stop after being trapped by a tree trunk or crown. Distinct colors for each individual indicate distinct MLLs. Despite displayed as a 2D-reproduction, all simulations have a ‘3D behavior’, once the model assigns distinct heights of each tree trunk and crown. Therefore, each tree could be thought of as a disk-shaped crown with a cylindrical trunk attached underneath its core. Moreover, despite that the representation shows a 2D-movement, each dispersed seed also changes its altitude at each movement step.

*5. Initialization and input data*

The simulation can start with (i) a representation of the empirical landscape; or with (ii) an artificial landscape. In the first case, 20 trees with specific heights of trunks and crowns are distributed over the area in specific positions (Table S1). In the second case, the set number of trees are distributed randomly over the available area with a previously set average crown area and hosting no *T. recurvata* individuals. All simulations are run in a homogeneous environment, where, beyond chosen wind speed, only the number of trees and their underlying crown overlapping affect the dispersal speed.

*6. Submodels*

In the following, details of the main processes are presented.

1. **Update**

Every time step (i.e. simulated month), the age and energy of all bromeliads are updated. The age is updated adding a month, and the energy is updated according to their energy budget that reduces the energy spent for metabolism from the energy produced through photosynthesis:

- Energy expense: the metabolism processes take a 0.3 and 0.5 unit of the accumulated energy of, respectively, adults (age > 12 months) and seedlings (age < 12 months). This difference is due to the greater energy spent by seedlings for their growth;
- Energy gain: photosynthesis adds a variable amount of energy, according to the following function:

(1)

energy gain = (((2-(1-periphery))/shading)/compet)

where *periphery* is where in tree crown the individual is located (individual_altitude_/tree_height_), *shading* is the number of tree crowns that cover the individual’s location, and *compet* is the number of other adult individuals in a radius of 15 cm from the focal individual.

2. **Reproductive season**

The reproductive season takes place every year (i.e. every 12 time steps representing one month each), where new individuals of *T. recurvata* are added from outside of the system (regional seed rain) or from other reproducing individuals in the system. All individuals in the model with at least 10 units of energy reproduce up to twice during their lifespan, producing new seeds and new ramets. Although *T. recurvata* be a monocarpic species (i.e. reproduce only once during their life cycle (Mercier & Endres 1999) it was an strategy to reduce the memory consumption of the model, since new shoots are produced just after the reproductive season, overlapping the older ramets. After reproduction each individual loses two thirds of its original energy.

*Regional seed rain*

A previous set number of regionally dispersed seeds are included in the simulation. Their alleles in each of the defined SSR loci, are randomly chosen between the shortest and longest alleles found in the empiric study, in accordance with their motifs (dinucleotide or trinucleotide) and with the heterozygosity observed in the empirical study. The dispersal origin of these seeds could be previously defined as north, south, east, west, or random. Their starting height are randomly chosen from 1 meter to up to five meters more than the average height of trees in the simulated landscape. The color which can be observed in the 2D-display for each seed is chosen randomly and can be seen as a new MLL entering into the simulated landscape. If an individual reproduces during the simulation, its color is inherited by their offspring.

The regional seed rain can also be increased linearly every year throughout the simulation. The increment can be linear:

seedrain^regional^ = seeds + (seeds * year)

(2)

where the parameter *seeds* is the previously set number of seeds in the regional seed rain. This increment mimics a growth of source populations outside the simulated landscape.

*Seed production*

The number of seeds that a reproducing individual produces is ten times its amount of energy. Each newly produced seed starts to move from the same location where it was produced in a random direction. As *T. recurvata* is a cleistogamous species, each produced seed has a genotype that is a random recombination of the alleles of its mother. However, if the mutation option in the model is ‘on’, each allele of the new seeds genotypes can gain or lose repeat units by chance (i.e. Stepwise Mutation Model), simulating the process known as “DNA replication slippage” of microsatellites, in a rate that is also previously set in a range between 10^-2^ and 10^-6^. Therefore, in this model, mutation is the single source of variation within MLLs. To track back MLLs, each new seed has a flag indicating the ID of the first individual of its MLL that arrived in the simulation.

*Clonal reproduction*

During the reproductive seasons, the same bromeliads that produced seeds also reproduce vegetatively, producing more ramets (hereafter referred as ‘clones’) according to their energy (N_clones_ = energy/5). Half of the initial energy of the reproducing bromeliad is shared equally among all new ramet. The clones stay attached close to their mother and have the same multi-locus genotype (MLG) and multi-locus lineage (MLL).

3. **Dispersal seed movement**

*move*

*T. recurvata* seeds are only dispersed by wind during the reproductive season (here delimited by a month within a year). Their movement is randomly divided into five ‘steps’, representing the distinct conditions they can face throughout the landscape. The wind speed is previously set by the user and is measured by an arbitrary metric that is related to the seed dispersal speed during the reproductive season. This metric represents how many model patches (0.01 m^2^) the seed moves at each step.

During a reproductive season, a dispersal seed moves straightforward, in a randomly chosen direction, losing altitude by chance. The distance the seed moves at each movement step depends on its position, above or below the average height of tree crowns in the surrounding ca. 3 meters. If the seed is above this threshold, its speed is five times the set wind speed and its altitude reduces 0.1 m at each movement step. However, if the seed is below this threshold, the wind speed effect on its movement decreases as the following function:

(3)

distance_step_ = speed_wind_ - (2 * shading - (0.1 * altitude))

where, *shading* is the number of canopies that covers the individual site and *altitude* is the location of the dispersed seed in the air, measured from the ground. According to the above function, the number of trees crowns covering a site reduces the speed of seed dispersal, but this effect is weakened by the seed’s altitude inside the canopy. In other words, the higher a seed is in the canopy, and the fewer trees are in the landscape, the greater is the effect of wind on its dispersal. Seeds below the canopy threshold also decrease in altitude 0.5 m at each movement step.

If a seed reaches to a tree trunk, it will be captured, but, if it reaches a tree crown, the model calculates its capture probability. This probability is previously set in the model by the user (ranging from 0-100%) and it is multiplied by the number of tree crowns that cover the space. All seeds that reach the ground or the landscape margins, or that are not captured during the reproductive season, die before the next month.

4. **Seed germination and seedling survival**

From all seeds that enter the simulated landscape or are produced in it, the model calculates the germination probability that is previously set by the user (from 0 to 100) and a 65 % of chance of seedling survival.

5. ***Mortality***

A bromeliad dies when its age reaches its lifespan limit of three years (i.e. 36 months) or when its energy is lower or equal to zero. To reach this energy value the energy gain of a bromeliad has to be lower than its expense. The energy of a bromeliad can also be reduced in a stochastic event, if it is activated in the model. If so, it can occur in a random month of each half time of the simulated period of 10 years. During these events all bromeliads are affected losing 5-25 energy units.

*7. References*

Benzing, D. H. (2000). *Bromeliaceae: profile of an adaptive radiation*. Cambridge: Cambridge University Press.

Benzing, D. H. (2012). *Air Plants: epiphytes and aerial gardens*. New York, NY: Cornell University Press.

Birge, W. I. (1911). The anatomy and some biological aspects of the “ball moss”, tillandsia recurvata L. *Bulletin of The University of Texas*, *194*(20).

Chaves, C. J. N., Aoki-Gonçalves, F., Leal, B. S. S., Rossatto, D. R., & Palma-Silva, C. (2018). Transferability of nuclear microsatellite markers to the atmospheric bromeliads Tillandsia recurvata and T. aeranthos (Bromeliaceae). *Brazilian Journal of Botany*, (August). doi:10.1007/s40415-018-0494-4

Chilpa-Galván, N., Márquez-Guzmán, J., Zotz, G., Echevarría-Machado, I., Andrade, J. L., Espadas-Manrique, C., & Reyes-García, C. (2018). Seed traits favouring dispersal and establishment of six epiphytic Tillandsia (Bromeliaceae) species. *Seed Science Research*, 1–11. doi:10.1017/S0960258518000247

Flores-Palacios, A., García-Franco, J. G., & Capistrán-Barradas, A. (2015). Biomass, phorophyte specificity and distribution of Tillandsia recurvata in a tropical semi-desert environment (Chihuahuan Desert, Mexico). *Plant Ecology and Evolution*, *148*(1), 68–75.

GBIF Secretariat (2017). GBIF Backbone Taxonomy. Checklist dataset https://doi.org/10.15468/39omei accessed via GBIF.org on 2018-10-23.

McWilliams, E. (1992). *Chronology of the natural range expansion of Tillandsia recurvata (Bromeliaceae) in Texas*. *Contributions to Botany* (Vol. 15).

Mercier, H., Endres, L. (1999) Alteration of hormonal levels in a rootless epiphytic bromeliad in different phenological phases. *Journal of Plant Growth Regulation*, 18, 121-125.

Nathan, R., Horn, H. S., Chave, J., & Levin, S. A. (2002). Mechanistic models for tree seed dispersal by wind in dense forests and open landscapes. In *Seed Dispersal and frugivory-Ecologie, Evolution, Conservation* (pp. 69–82). doi:10.1079/9780851995250.0069

Orozco-Ibarrola, O. A., Flores-Hernández, P. S., Victoriano-Romero, E., Corona-López, A. M., & Flores-Palacios, A. (2015). Are breeding system and florivory associated with the abundance of Tillandsia species (Bromeliaceae)? *Botanical Journal of the Linnean Society*, *177*(1), 50–65. doi:10.1111/boj.12225

Smith, A. K., Martin, C. E., & Lüttge, U. (1989). Gas exchange and water vapor uptake in the atmospheric CAM bromeliad Tillandsia recurvata L.: the influence of trichomes. *Botanica Acta*, *102*, 80–84.

Smith, L. B., & Downs, R. J. (1977). Tillandsioideae (Bromeliaceae). In *Flora Neotropica* (pp. 661–1178).

Soltis, D. E., Gilmartin, A. J., Rieseberg, L., & Gardner, S. (1987). Genetic Variation in the Epiphytes Tillandsia ionantha and T. recurvata (Bromeliaceae). *American Journal of Botany*, *74*(4), 531–537.

Wilensky, U. (1999). NetLogo. http://ccl.northwestern.edu/netlogo/. Center for Connected Learning and Computer-Based Modeling, Northwestern University, Evanston, IL.
